# Reconciling fMRI and EEG indices of attentional modulations in human visual cortex

**DOI:** 10.1101/391193

**Authors:** Sirawaj Itthipuripat, Thomas C Sprague, John T Serences

## Abstract

Functional magnetic resonance imaging (fMRI) and electroencephalography (EEG) are the two most popular non-invasive methods used to study the neural mechanisms underlying human cognition. These approaches are considered complementary: fMRI has higher spatial resolution but sluggish temporal resolution, whereas EEG has millisecond temporal resolution, but only at a broad spatial scale. Beyond the obvious fact that fMRI measures properties of blood and EEG measures changes in electric fields, many foundational studies assume that, aside from differences in spatial and temporal precision, these two methods index the same underlying neural modulations. We tested this assumption by using EEG and fMRI to measure attentional modulations of neural responses to stimuli of different visual contrasts. We found that equivalent experiments performed using fMRI and EEG on the same participants revealed remarkably different patterns of attentional modulations: event-related fMRI responses provided evidence for an additive increase in responses across all contrasts equally, whereas early stimulus-evoked event-related potentials (ERPs) showed larger modulations with increasing stimulus contrast and only a later negative-going ERP and low-frequency oscillatory EEG signals showed effects similar to fMRI. These results demonstrate that there is not a one-to-one correspondence between the physiological mechanisms that give rise to modulations of fMRI responses and the most commonly used ERP markers, and that the typical approach of employing fMRI and EEG to gain complementary information about localization and temporal dynamics is over-simplified. Instead, fMRI and EEG index different physiological modulations and their joint application affords synergistic insights into the neural mechanisms supporting human cognition.

## Introduction

For decades, a growing number of studies have employed fMRI and EEG as complementary methods to study neural mechanisms underlying human cognition and there is a widespread notion that any modulations in stimulus-evoked responses measured using functional magnetic resonance imaging (fMRI) or via electroencephalography (EEG) merely reflects different assays of the same underlying neural activity changes. For example, there is a long tradition in the study of selective attention to use both fMRI and EEG to assess attention-induced gain amplification of sensory signals, with the former measure used for spatial localization and the later for tracking precise timing (Hillyard and Anllo-Vento, 1998; Mangun et al., 1998; Martinez et al., 1999; Martínez et al., 2001; Noesselt et al., 2002; Di Russo et al., 2002, 2005, 2007; Busse et al., 2005; Novitskiy et al., 2011; Zhang et al., 2012; Di Russo and Pitzalis, 2013; Chen et al., 2014; Green et al., 2017). In one seminal line of work, for example, fMRI was used to localize the likely sites of attentional modulations in human visual cortex and then EEG, coupled with source localization procedures, was used to examine the temporal profile of attentional modulations from the fMRI-identified ‘seed’ regions (Hillyard and Anllo-Vento, 1998; Martinez et al., 1999; Martínez et al., 2001; Di Russo et al., 2002, 2005, 2007; Di Russo and Pitzalis, 2013).

Clearly, fMRI and EEG are different assays of neural activity, with fMRI measuring changes in blood volume and the ratio of oxygenated to deoxygenated hemoglobin (Logothetis et al., 2001; Logothetis, 2002, 2008) and EEG measuring electrical potentials on the scalp generated by coherent activity in large populations of cortical neurons (Luck, 2012; Lopes da Silva, 2013). That said, beyond this difference in the source of the signals, most studies implicitly assume that modulations in both measures are proxies for the same underlying changes in neural activity. However, there are many hints in the literature that fMRI and EEG are not simply two complementary means of assaying the same neural modulations. For example, when neural responses are measured as a function of stimulus contrast to obtain contrast response functions (CRFs), different types of attentional modulations have been observed across techniques. Results from fMRI unanimously support a mechanism whereby attention increases the evoked response to all stimuli equally, regardless of their contrast, while results from EEG support a contrast-dependent response modulation (either a horizontal shift of the CRF, called ‘contrast gain’, or a multiplicative scaling of the CRF, called ‘response gain’ (Figure 1a; reviewed in (Lee and Maunsell, 2009; Reynolds and Heeger, 2009; Hara et al., 2014; Itthipuripat and Serences, 2015)). While these different types of attention effects could be due to differences in task designs, stimulus properties, recording sites, training duration, and subjects’ attentional strategy and expertise (Reynolds and Heeger, 2009; Herrmann et al., 2010; Itthipuripat et al., 2014b, 2017; Zhang et al., 2016), another reasonable source of divergence is the neural response properties to which each measurement is most sensitive (Boynton, 2011; Itthipuripat and Serences, 2015).

**Figure 1.**
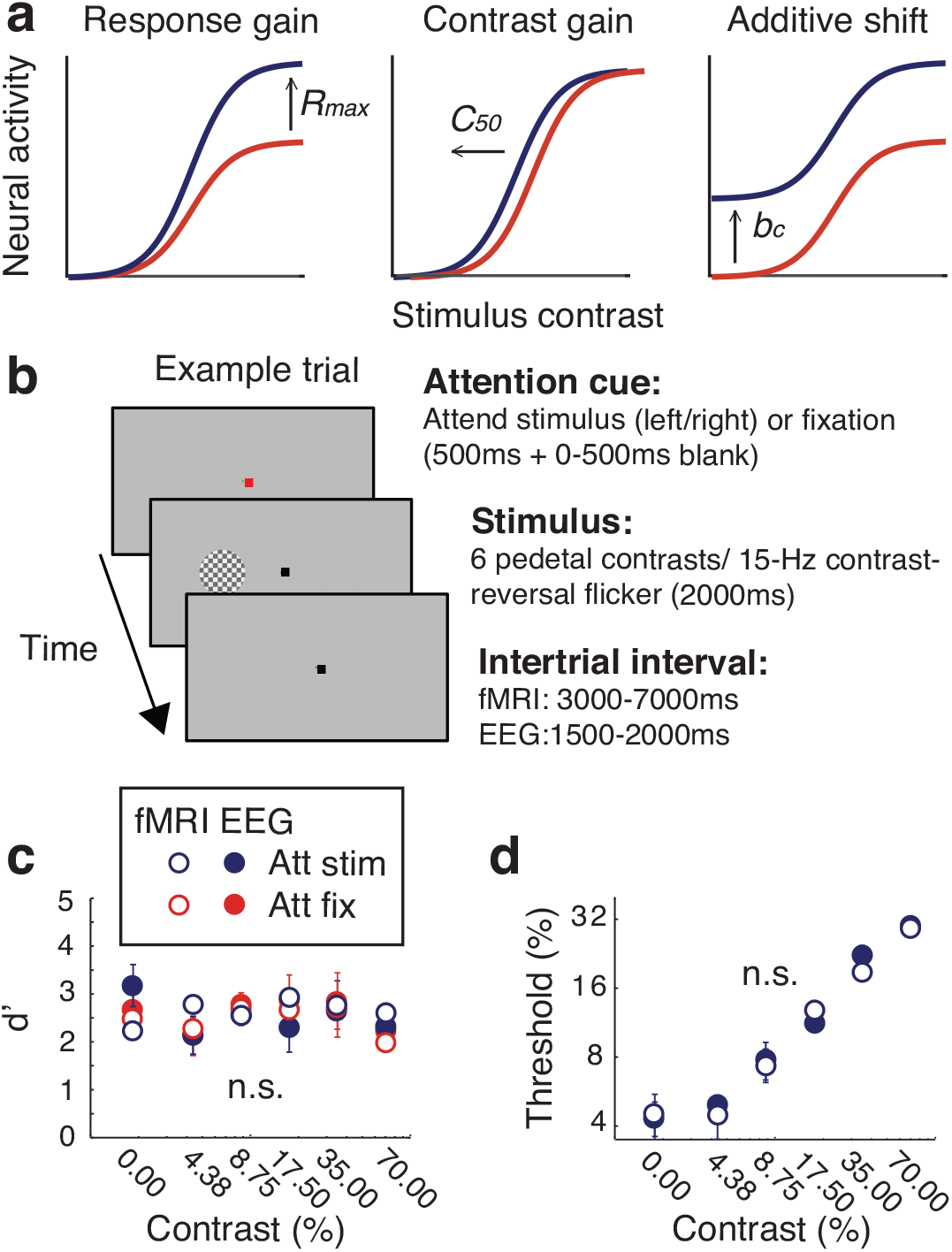
Predictions, behavioral task, and behavioral results. **(a)** Different patterns of attentional gain modulations in contrast response functions (CRFs) measured in visual cortex. Past studies suggest that attention induces changes in responses gain (*R_max_*), contrast gain (*C_50_*), or the CRF baseline activity (*b_c_*) depending on the size of the spatial scope of attention, stimulus properties, and training duration (Reynolds and Heeger, 2009; Herrmann et al., 2010; Itthipuripat et al., 2014b, 2017). Here, we predicted that measurement modality (e.g., fMRI vs. EEG) was another source of differences in these modulatory patterns between studies. **(b)** The spatial attention task. Each trial started with a color cue instructing human participants to attend to the central fixation point (i.e., attention-fixation) or to covertly attend to a stimulus on the left or the right of the fixation (i.e., attend-stimulus). The stimulus was flickered at 15 Hz (contrast-reversing) and its contrast was pseudo-randomly and uniformly drawn from one of 6 possible values (0, 4.38, 8.75, 17.50, 35, and 70% contrast values). Participants (*n* = 7) detected contrast changes at fixation or at the stimulus location in the attention-fixation and the attend-stimulus conditions, respectively (25% target trials). **(c)** Behavioral performance was equated across all contrast levels, attention conditions, and measurement modalities. **(d)** Contrast thresholds in the attend-stimulus condition were not different across measurement modalities. Error bars represent within-subject S.E.M.

Here, we tested the hypothesis that a major source of discrepancies between results from different measurement techniques is directly attributable to differences in their *physiological* resolution, and thus the types of neural modulations that drive changes in the signals being recorded. So far, it is difficult to draw any solid conclusion by comparing results across existing studies that used different neural measurements because different neural metrics are often measured from different individuals performing different behavioral tasks with different levels of difficulty— factors that have been shown to alter neural patterns of attentional modulations (Lee and Maunsell, 2009; Reynolds and Heeger, 2009; Hara et al., 2014; Itthipuripat and Serences, 2015). Furthermore, even those studies that did make fMRI and EEG measurements within the same participants, the data were not quantified in such a way as to reveal qualitative differences and to critically evaluate the assumption that the same neural generators gave rise to modulations in both measures (Hillyard and Anllo-Vento, 1998; Mangun et al., 1998; Martinez et al., 1999; Martinez et al., 2001; Noesselt et al., 2002; Di Russo et al., 2002, 2005, 2007; Busse et al., 2005; Novitskiy et al., 2011; Zhang et al., 2012; Di Russo and Pitzalis, 2013; Chen et al., 2014; Green et al., 2017).

## Materials and Methods

### Participants

Seven neurologically healthy human observers (19-32 years old, 3 females, 1 left-handed) wih normal or corrected-to-normal vision were recruited from the University of California, San Diego (UCSD) community. All participants provided written informed consent, approved by the human subjects Institutional Review Board at UCSD and the experiment was conducted under the protocol that followed the Declaration of Helsinki. The participants were compensated for $10, $15, and $20 per hour for participating in behavioral training, EEG, and fMRI recording sessions, respectively. Two of the participants are authors (S.I and T.C.S.) and were not compensated.

### Stimulus presentation

During behavioral and EEG recording sessions, stimuli were presented on a PC running Windows XP using MATLAB (Mathworks Inc., Natick, MA) and the Psychophysics Toolbox (version 3.0.8). Participants sat 60 cm from the CRT monitor (60 Hz refresh rate) in a sound-attenuated and electromagnetically shielded room (ETS Lindgren). During fMRI scanning sessions, we presented stimuli using a contrast-linearized LCD projector (60 Hz) on a rear-projection screen mounted at the foot of the scanner (110 cm wide, ~4.1 m viewing distance). All stimuli appeared on a neutral gray background.

### fMRI and EEG main tasks

Throughout the main EEG task, we instructed participants to fixate on the dark gray fixation point located at the center of the gray screen. Individual trials started with a 500 ms color cue, instructing participants either to covertly attend to the checkerboard stimulus (radius = 1.45 ^o^ visual angle) located 3.6° to the left (a red cue) or right (a blue cue) relative to fixation (attend-stimulus condition). Alternatively, subjects were instructed to maintain fixation on the gray fixation dot while ignoring the peripheral stimulus (indicated with a green cue; attend-fixation condition). The checkerboard stimulus appeared 500-1000 ms after cue onset and the stimulus continued flickering at 15 Hz (contrast reversal) for 2000 ms. On each trial, the checkerboard stimulus had one of the following Michelson contrast levels: 0%, 4.375%, 8.75%, 17.5%, 35%, and 70% (logarithmically-spaced). In fMRI sessions, the inter-trial interval (ITI) varied from 3000-7000 ms, and in EEG experiments, it varied from 1500-2000 ms. On 25% of the attend-stimulus trials, the stimulus contained a constant contrast increment (target trials). On 25% of the attend-fixation trials, the grey fixation dot contained a constant contrast increment (target trials). The contrast increment in both attend-stimulus and attend-fixation target trials appeared anytime from 600 ms to 1300 ms after the stimulus onset. Subjects were instructed to press a button with their right index finger as quickly and accurately as possible when they saw this contrast increment. The contrast increment of the fixation dot and the contrast increment of the checkerboard stimulus was separately determined for each pedestal contrast level on a block-by-block basis to fix the hit rate at ~75% across all stimulus and task conditions.

Each subject completed 1 fMRI session and 2 EEG sessions with each session completed on different days. The fMRI experiment contained 6 blocks of trials, while the EEG experiment contained 20 of blocks of trials in total. Every 2 blocks of the main task contained 96 trials in total where all stimulus and attention conditions were counterbalanced: 2 attention conditions (attend-stimulus/ attend-fixation) x 2 stimulus location (left/right hemifield) x 6 pedestal contrast x 4 repeats. The order of stimulus and attention conditions was pseudorandomized within these 2-block sequences. To control for any possible effects of learning that might occur across sessions, 4 participants first underwent 2 EEG sessions followed by an fMRI session while the other 3 participants first underwent an fMRI session followed by 2 EEG sessions. We obtained fewer trials of fMRI data than EEG data because the observed fMRI signals had a relatively high signal-to-noise ratio.

Before participants began the first recording session (either EEG or fMRI), they underwent a 2.5-hour behavioral training session on an identical task, except that there were targets on 50% of the trials instead of 25%, there was no response deadline, and subjects had to answer whether each trial was a target (a stimulus with a contrast increment) or a non-target trial by pressing one of the two corresponding buttons on a keyboard. During this training session, the contrast thresholds were estimated using a staircase procedure that was applied independently for each attention condition and each pedestal contrast level. Three successive correct responses (either a hit or a correct rejection) led to a 0.5% decrease in the Δc that defined the target stimulus, while one incorrect response led to a 0.5% increase in Δc (either a miss or a false alarm). Trials from the first five reversals were excluded and the mean values of the contrast increments from remaining trials were used as contrast detection thresholds in the first block of the first EEG or fMRI recording session.

### Behavioral Analysis

We first computed perceptual sensitivity or behavioral d-prime (d’), using the following equation: d’ = Z(hit rate) - Z(false alarm rate), where Z is the inverse of the cumulative distribution function of the Gaussian distribution. To test if d’ were equated across contrast levels (6 levels), attention conditions (attend-stimulus/attend-fixation), and measurement modalities (fMRI/EEG), we used a 3-way repeated-measured ANOVA with these independent measures as with-in subject factors. In addition, we also used a separate 2-way repeated-measures ANOVA to test the main effects of contrast and measurement modality and their interactions on the behavioral contrast detection thresholds.

### fMRI functional localizers

Participants performed 1-2 blocks of a functional localizer task to identify voxels that were visually responsive to the portion of the visual field subtended by the stimulus in the main task. Subjects maintained fixation while ignoring the localizer stimulus. The peripheral checkerboard stimulus was 100% contrast and presented at the same size and location as in the main task. It flickered at 15 Hz and alternately appeared in the left and right stimulus locations for 8 s/trial. Subjects responded with a button press when they perceived a brief and small contrast change at the fixation point; contrast detection targets could appear between 2-3 times per 8 s trial.

To estimate the spatial sensitivity profile of each voxel during the “training” phase of the IEM analysis (see below), participants performed 7-8 blocks of a spatial mapping task, with all participants performing 4 blocks using low-contrast mapping stimuli (50% contrast), and one participant performing 3 blocks using high-contrast mapping stimuli (100% contrast), while the remaining participants performed 4 high-contrast mapping blocks. To ensure IEMs for all participants were estimated with an equivalent amount of data, we used the low-contrast mapping data for all analyses reported here. On each trial, a 15 Hz flickering checkerboard stimulus 2.90° in diameter appeared at a different location on the screen, selected from a 8 x 4 square grid (1.45° horizontal/vertical spacing) and jittered on each trial (+-0.725° in X,Y independently). We also included 6 null trials in which no checkerboard stimulus appeared. On all trials (stimulus-present and null trials), participants carefully monitored for a brief dimming of the fixation point, which acted as a target stimulus (1 target per trial; targets appeared on 50% of trials), and participants responded with a button press when the dimming occurred.

### fMRI retinotopic mapping procedure

Striate and extra-striate visual areas (V1, V2v, V3v, V2d, V3d, hV4) were defined by standard retinotopic mapping procedures, using a rotating counter-phase flickering checkerboard in conjunction with bowtie stimuli subtending the vertical and horizontal visual field meridians on alternating blocks, during separate scanner runs. The data were projected onto a computationally inflated gray/white matter boundary surface reconstruction for visualization (Engel et al., 1994; Sereno et al., 1995). V2 and V3 were combined by concatenating voxels so that any slight errors in drawing the horizontal meridian boundaries that separate them would not bias the inclusion of the localizer-defined voxels into one region or the other. This was especially important because the visual stimuli in this study were presented along the horizontal meridian, so imperfections in ROI definitions could result in erroneous conclusions about differences between V2 and V3, which we do not believe are possible to fairly assay with this stimulus setup. For the V1-hV4 ROI, we concatenated all voxels from all ROIs (V1, V2/V3, and hV4).

### fMRI Acquisition, Preprocessing and Analysis

We acquired fMRI data on a 3 Tesla research-dedicated GE MR750 located at the Keck Center for Functional MRI at the University of California, San Diego. We scanned all participants twice: once for a retinotopic mapping session, and once for the main task session (each scan ~2 hrs). During each session we acquired a high-resolution whole-head anatomical image used to align to the retinotopic mapping session (T1-weighted fast spoiled gradient echo sequence, 25.6 cm x 25.6 cm FOV, 256 x 192 acquisition matrix, 8.136/3.172 ms TR/TE, 192 slices, 9° flip angle, 1 mm isotropic voxel size).

We acquired task data using a Nova 32-channel head coil (Nova Medical) at 3 mm isotropic resolution using axial slices spanning occipital cortex, and also including parietal and frontal cortex (TR=2000 ms, TE = 30 ms, flip angle = 90°, 35 interleaved slices, 3 mm thickness, 0 mm gap, 19.2 x 19.2 cm FOV, 64 x 64 acquisition matrix). We acquired 179 volumes of data per run for the main task, 91 volumes for the IEM mapping task, and 118 volumes for the stimulus localizer task.

Preprocessing included unwarping using custom scripts implementing procedures from AFNI and FSL. All subsequent preprocessing occurred in BrainVoyager 2.6.1, including slice time correction, six-parameter rigid-body motion correction, high-pass temporal filtering to remove slow signal drifts over the course of each run, and transformation of data into aligned Talairach space. Then, the BOLD signal was normalized within each voxel for each run separately to Z-scores. All other analyses involved custom MATLAB scripts.

For fMRI analyses, we extracted the signal at each voxel on each trial using a GLM framework. We modeled each trial independently for the IEM mapping task and main contrast discrimination task (HRF: two-gamma, time-to-peak 5 s, undershoot peak at 15 s, response undershoot ratio 6, response and undershoot dispersion of 1). For the stimulus localizer task, we modeled all trials using a ‘left’ and a ‘right’ regresssor. For univariate analyses, we extracted activation from voxels significantly activated by the localizer task (q = 0.05, whole-brain FDR corrected), averaged across voxels responsive to the left or right stimulus, and sorted trials by contrast and attention condition.

For multivariate analyses, we modeled the response of each voxel as a linear combination of a discrete set of spatial filters, or ‘information channels’ (see (Sprague et al., 2016) for a detailed description of the analysis framework). We modeled channels as a rectangular grid, 9 x 5, of 1.81° full-width half-maximum (FWHM) round filters, spaced by 1.449° horizontally/vertically:

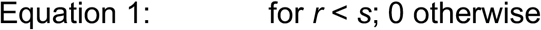

Where *r* is the distance from the filter center and s is a “size constant” reflecting the distance from the center of each spatial filter at which the filter returns to 0. Values greater than this are set to 0, resulting in a single smooth round filter at each position along the triangular grid (*s* = 4.554°).

This rectangular grid of filters forms the set of information channels and each mapping task stimulus is converted from a contrast mask (1’s for each pixel subtended by the stimulus, 0’s elsewhere) to a set of filter activation levels by taking the dot product of the vectorized stimulus mask and the sensitivity profile of each filter. Once all filter activation levels are estimated, we normalize so that the maximum filter activation is 1.

Following previous reports (Brouwer and Heeger, 2009a; Sprague and Serences, 2013a), we model the response in each voxel as a weighted sum of filter responses:

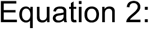

Where *B_1_* (*n* trials × *m* voxels) is the observed BOLD activation level of each voxel during the spatial mapping task (beta weight estimated from single-trial GLM), *C_1_* (*n* trials × *k* channels) is the modeled response of each spatial filter, or information channel, on each non-target trial of the mapping task (normalized from 0 to 1 across all channels and trials), and *W* is a weight matrix (*k* channels × *m* voxels) quantifying the contribution of each information channel to each voxel. Because we have more stimulus positions than modeled information channels, we can solve for *W* using ordinary least-squares linear regression:

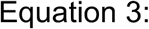

This step is univariate and can be computed for each voxel in a region independently. Next, we used all estimated voxel encoding models within a ROI () and a novel pattern of activation from the visual attention task (beta weight for each voxel estimated from single-trial GLM) to compute an estimate of the activation of each channel (, *n* trials × *k* channels) which gave rise to that observed activation pattern across all voxels within that ROI (*B_2_, n* trials × *m* voxels):

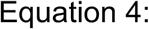

Once channel activation patterns are computed (Equation 4), we compute spatial reconstructions by weighting each filter’s spatial profile by the corresponding channel’s reconstructed activation level and summing all weighted filters together. This step aids in visualization, quantification, and coregistration of trials across stimulus positions, but does not confer additional information. To visualize these responses, we multiplied each channel’s filter profile by its activation measured during the task, and horizontally flipped trials in which the stimulus appeared on the left to align reconstructions as though all trials consisted of right stimuli.

Finally, to quantify these reconstructed stimulus representations, we fit a smooth surface to averaged reconstructions at each contrast and attention condition, within each ROI and participant:

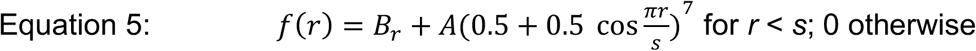

We implemented a coarse-to-fine fitting procedure, in which we first sampled a grid spanning center position −3 to 6° (0.33° spacing) horizontally (*x*), −3 to 3° (0.33° spacing) vertically (*y*), and FWHM (scaled version of *s*) from 0.25 to 22.25° (0.5° spacing), and fit amplitude (*A*) and reconstruction offset (*B_r_*) to a surface generated by the parameters at each point in the grid using least-squares regression. Then, we used the best-fit seed values from this initial coarse grid, defined by lowest RMSE, to seed a constrained optimization procedure to optimize for lowest RMSE. The constraints on *x, y*, and FWHM were identical to those spanned by the grid, and amplitude and baseline were additionally restricted to the range of −5:10 each. This resulted in one best-fit surface, parameterized by its amplitude (*A*), size (σ or FWHM), and reconstruction offset (*B_r_*) for each contrast and attention condition within each ROI for every participant. As described below, the amplitude parameter from this analysis was subjected to a second set of analyses to infer the contrast response function of each ROI for each attention condition.

### EEG recording, preprocessing and analysis

We recorded EEG data at a sampling rate of 512 Hz with a 64+8 electrode Biosemi ActiveTwo system (Biosemi Instrumentation), and placed two reference electrodes on the left and right mastoids. We also monitored blinks and vertical eye movements with four external electrodes placed above and below the eyes and horizontal eye movements with another pair of external electrodes placed near the outer canthi of the left and right eyes. The data were referenced online to the CMS-DRL electrode and the data offsets in all electrodes were maintained below 20 μV (a standard criterion for this active electrode system).

We preprocessed and analyzed EEG data using a combination of EEGlab11.0.3.1b and custom MATLAB scripts. We first re-referenced the continuous EEG data to the average of the EEG recorded from two mastoid electrodes. Then, we applied 0.25 Hz high-pass and 55 Hz low-pass Butterworth filters (3rd order) and segmented the data into epochs extending from −2500 ms before to 2500 ms after the stimulus onset. Artifact rejection was performed off-line by discarding epochs contaminated by eye blinks and vertical eye movements (more than ±80-150 μV deviation from zero; exact thresholds were determined on a subject-by-subject basis due to differences in the amplitudes of eye blink and vertical eye movement artifacts), horizontal eye movements (more than ±75 μV deviation from zero), excessive muscle activity, or drifts using threshold rejection and visual inspection on trial-by-trial basis, resulting in the removal of 12.61% (SD = 6.63%) of trials across subjects.

To obtain steady-state visually evoked potentials (SSVEPs), Fourier coefficients were calculated at 30 Hz (the second harmonic of the contrast-reversal flicker frequency of 15 Hz) and surrounding frequencies over the 2-s stimulus interval (0.5-256 Hz in consecutive 0.5 Hz steps). Next, the absolute values of the Fourier coefficients averaged across all artifact-free trials were computed separately for each attention condition (attend-stimulus/ attend-fixation), each stimulus location (left/right), each stimulus contrast level (0%-70%), and each electrode. The signal-to-noise ratios (SNR) of the SSVEP response for each stimulus contrast level and attention condition were calculated by dividing the amplitude of the second harmonic of the stimulus frequency (30 Hz) by the mean amplitude in the two frequency bins above and below the center frequency of 30 Hz (28.5-29Hz and 31-31.5 Hz respectively). We adopted this SNR metric following previous SSVEP studies to ensure that the modulations of the SSVEP were not confounded by any changes in broadband power at beta frequencies (Ding et al., 2006; Kim and Verghese, 2012; Verghese et al., 2012; Garcia et al., 2013; Itthipuripat et al., 2014b). We rearranged the SSVEP data so that electrodes ipsilateral and contralateral to the stimulus are positioned on the left and the right of the topographical map, respectively. We then collapsed the data obtained when the stimulus was presented in the left and the right hemifields. Finally, we plotted the SSVEP signals as a function of stimulus contrast to obtain the neural CRFs based on the SSVEP responses. We focused our SSVEP analysis on three posterior-occipital electrodes where the SSVEP SNR, averaged across all contrast levels, attention conditions, and participants was maximal.

To obtain event-related potentials (ERPs), we baseline corrected from −200-0 ms relative to the stimulus onset and then computed the algebraic mean of the EEG data previously sorted into different contrast and attention conditions. The ERP data were also rearranged so that electrodes ipsilateral and contralateral to the stimulus are positioned on the left and the right of the topographical map, respectively, and we collapsed the data obtained when the stimulus was presented in the left and the right hemifields. We focused our ERP analysis on three ERP components, including the visual P1 component from 110-130 ms at the posterior occipital electrodes, the visual N1 component from 150-170 ms at the contralateral posterior occipital electrodes, the late positive deflection (LPD or P3) from 250-350 ms at the midline posterior electrodes, and the contralateral late negative-going wave (CLN) from 800-2000 ms at the posterior occipital electrodes. The electrodes of interest for each of these ERP components were different sets of three electrodes that showed the maximal response amplitude averaged across all contrast levels, attention conditions, and participants. Similar to the analysis of SSVEP responses, we plotted the amplitude of these ERP components from the electrodes of interest as a function of stimulus contrast to obtain the neural CRFs.

Last, we examined post-stimulus changes in posterior alpha activity. To do so, we wavelet-filtered the artifact-free epoched EEG data using a Gaussian filter centered at 10-12 Hz with a time-domain standard deviation ranging from 83-100 ms (see similar methods in (Canolty et al., 2007, 2009; Itthipuripat et al., 2013)). Next, we computed changes in the alpha amplitude during the 2 s stimulus duration relative to the pre-cue period by subtracting out the mean alpha amplitude averaged across −500-0 ms before the cue onset. Finally, we plotted post-stimulus alpha amplitude as a function of stimulus contrast to obtain the neural CRFs, and we focused our alpha analysis on three contralateral posterior-occipital electrodes where the reduction in alpha amplitude, averaged across all contrast levels, attention, and subjects conditions, was maximal.

### Quantifying contrast response functions

To examine whether attention induces either response gain, contrast gain, or baseline shifting in the CRF as measured using fMRI and EEG measurements (see details above), we fit a Naka-Rushton function to the data as follows. First, we used a bootstrapping procedure to resample subjects with replacement and we computed the averaged response for each contrast level and each attention condition across the resampled subject labels. Then we fit the resampled data (12 data points: 2 attention conditions x 6 contrast levels) with the following Naka-Rushton equation:

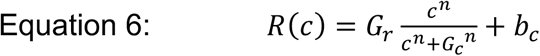

The fitting procedure was performed with 8 free parameters: 2 response gain factors (*G_r_*), 2 contrast gain factors (*G_c_*), 2 baseline parameters (*b_c_*), and 2 exponents (*n*); one for each attention condition (attend-stimulus and attend-fixation). We used the MATLAB function “fmincon” to minimize the root mean squared error between the data and the fit function, under a set of constraints. For fMRI fits, *G_r_* was restricted to be positive, with a maximum of 5 BOLD Z-score units; *G_c_* was restricted within the range of 0 to 100 (% contrast), and CRF baseline activity was restricted to an absolute value of 3 BOLD Z-score units. For EEG fits, *G_r_* was restricted to be within a range of −20 to 20 μV; *G_c_* was restricted within the range of 0 to 100 (% contrast), and *b_c_* was restricted to be within a range of −6 to 6 μV. For both EEG and fMRI fits, the exponent *n* was restricted within 0.1 and 5.

Since the *G_r_* and *G_c_* parameters control the response and contrast gain of the function where the contrast axis ranges from zero to ∞, the *G_r_* and *G_c_* parameters could in principle exceed the realistic range of stimulus contrast (0-100% contrast). Thus, instead of directly comparing *G_r_* and *G_c_* parameters across conditions we obtained the maximum response (*R_max_*) by evaluating the Naka-Rushton equation at *c* = 100%, and the semi-saturation parameters (the contrast value at half the *R_max_, C_50_*) as they respectively capture response gain and contrast gain of the CRFs over the realistic range of stimulus contrast values (see similar methods in (Itthipuripat et al., 2014b)). For fMRI data, we fit either the univariate mean BOLD response contralateral to the stimulus position (Figures 2a; S1), or the amplitude of the best-fit surface to the IEM-based image reconstructions (Figures 2d; S2). For EEG data, this fitting procedure was done separately for each of the 64 EEG electrodes.

**Figure 2.**
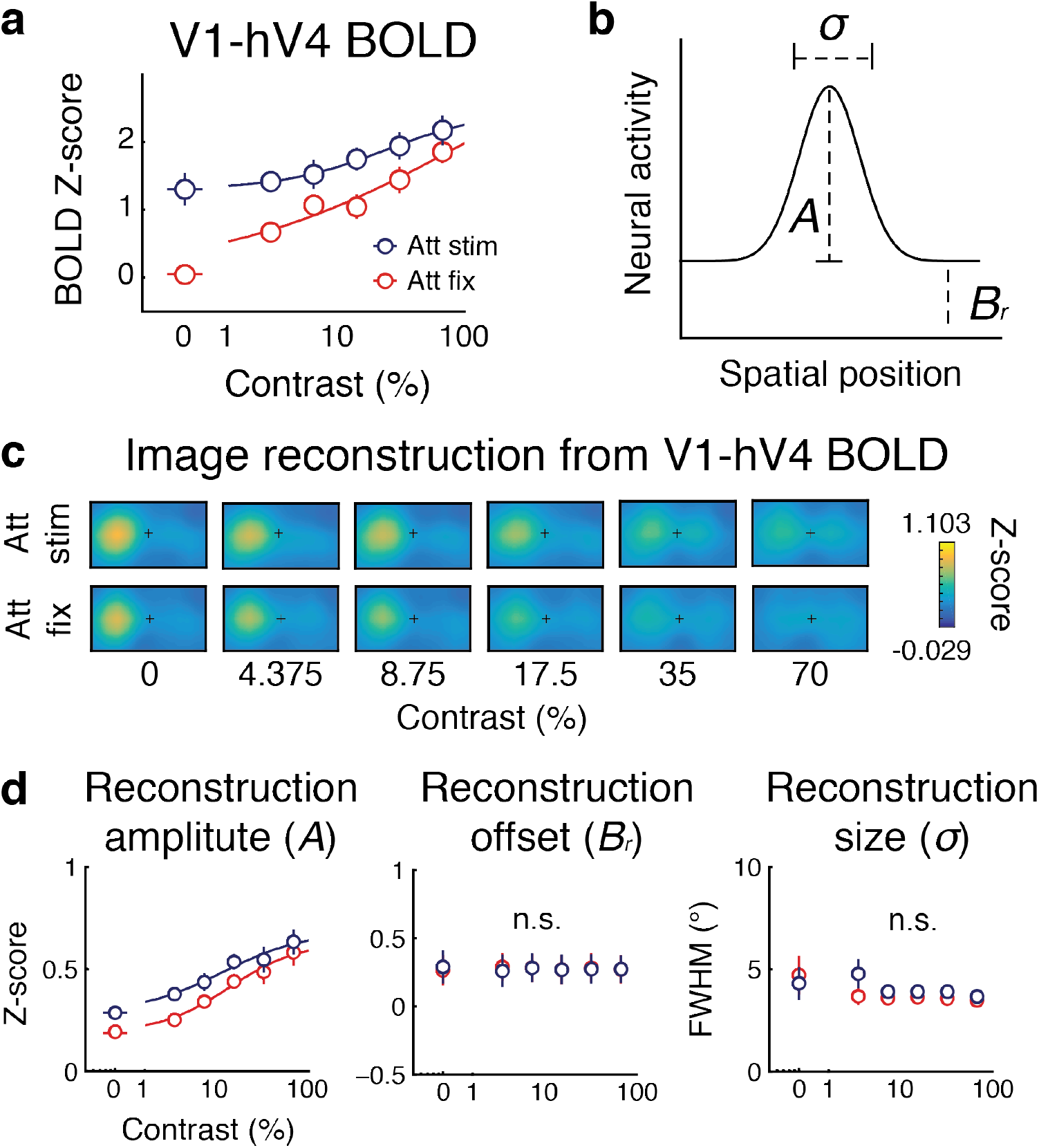
Spatial attention increases baseline CRF baseline activity as measured with fMRI. **(a)** Spatial attention altered the CRF baseline activity (*b_c_*) based on univariate fMRI activity measured in human visual areas V1-hV4. **(b)** A Gaussian-like function used to fit and describe the spatial reconstruction of fMRI activity. **(c)** fMRI-based image reconstructions of stimuli of different contrast levels across different attention conditions. Trials are aligned such that stimuli were presented on the left of these reconstructions. **(c)** Attention and contrast effects on the reconstruction amplitude (*A*), size (*σ* or FWHM), and reconstruction offset (*B_r_*) parameters. Similar to the univariate fMRI result, spatial attention altered the CRF baseline activity (*b_c_*) of reconstruction amplitude (*A*). There was no attention or contrast effect or an interaction between the two factors on the other reconstruction parameters. Error bars represent 68% CIs from resampling procedures. Att stim: attend stimulus; Att fix: attend fixation.

To obtain bootstrapped distributions of *R_max_, C_50_, b_c_*, and *n* parameters, this resampling and refitting procedure was performed 10,000 times, with each iteration resampling over participants with replacement. To test the significance of the attention effect on each of these parameters, we compiled the bootstrapped distribution of the differences between the estimated fit parameters in the attend-stimulus and attend-fixation conditions and computed the percentage of values in the tail of this distribution that were less- or greater than zero. We used two-tailed statistical tests throughout to be conservative, so we doubled this proportion to obtain each *p*-value.

### Data/software availability

All data and analysis code supporting reported results is available in the author’s Open Science Framework and GitHub repositories (upon publication).

## Results

### Behavioral results

To test the hypothesis that fMRI and EEG index fundamentally different aspects of neural activity, we employed both techniques to measure attentional modulations of neural CRFs in the same human subjects performing the same visual spatial attention task under matched stimulus conditions and difficulty levels (Figure 1b). Across fMRI and EEG recording sessions, seven participants detected a rare incremental change in the contrast of a target (25% of all trials) at the fixation point or at the stimulus location (left or right of fixation; the circular checkerboard stimulus flickered at 15 Hz contrast reversal) following a central color cue. As shown in Figure 1c, behavioral perceptual sensitivity (d’) was equated across stimulus contrast levels (0-70% pedestal contrast, equally spaced on a logarithmic scale), attention conditions (attend-fixation vs. attend-stimulus), and measurement modalities (fMRI vs. EEG). Thus, there was no main effect of stimulus contrast (F(5, 30) = 2.46, p = 0.056), attention (F(1, 6) = 0.16, p = 0.70), or measurement modality (F(1, 6) = 0.00, p = 0.96), nor an interaction between any combination of these three factors (F(5, 30) = 0.67, F(5, 30) = 1.40, F(1, 6) = 0.36, and F(5, 30) = 0.91 for the interactions between contrast and attention, between contrast and modality, between attention and modality, and between all three factors, respectively, with all p’s ≥ 0.25). The contrast detection thresholds increased as a function of stimulus contrast consistent with many past studies (a significant main effect of contrast: F(5, 30) = 233.40, p <0.001) (Legge and Foley, 1980; Ross et al., 1993; Boynton et al., 1999; Gorea and Sagi, 2001; Huang and Dobkins, 2005; Pestilli et al., 2011; Itthipuripat et al., 2014a, 2017). Moreover, behavioral performance did not differ significantly across fMRI and EEG sessions (Figure 1d) (no main effect of measurement modality and no interaction between contrast and measurement modality: F(1, 6)=0.61 and F(5, 30) = 2.46 with p = 0.464 and p = 0.055, respectively). Overall, the similarity of behavioral results across measurement modalities ensured that the difference in attentional modulations measured via fMRI and EEG were not be due to any difference in factors such as task difficulty or strategy.

### Univariate fMRI results

Univariate fMRI analyses qualitatively showed that spatial attention induced an additive shift in the fMRI-based CRFs (i.e., attention increases BOLD activity; Figure 2a and Figure S1). To quantify the shape of the CRFs and their modulation with attention, we fit a standard Naka-Rushton equation that has parameters for the maximum response (*R_max_*), the point at which the response reaches 50% of max (*C_50_*), and the CRF baseline activity or y-intercept of the CRF (*b_c_*, see Online Methods). Across all contralateral retinotopic early visual ROIs V1, V2/V3 and hV4, attention reliably increased the CRF baseline activity (*b_c_*, resampling tests described in Online Methods, all p’s ≤ 0.001 across all visual areas, also see Table S1). In addition, in these regions we observed a significant attention-induced reduction in response gain or *R_max_* (the difference between responses at 0% and 100% contrast; resampling tests, all p’s < 0.001). There were less consistent effects of attention on the other parameters of the CRFs across different visual areas: *p*-values ranged from 0.026-0.914 for *C_50_* (the contrast at which the best fitting CRF reached half its response at 100% contrast) and *p*-values ranged from 0.025-0.112 for the exponent *n* (the steepness of the fit CRF; Figure S1c, resampling tests, see Table 1 for all statistical results).

### Multivariate fMRI results

Next, to further explore the impact of attention on the information content of fMRI-based on activation patterns across entire visual areas rather than their mean activation level, we reconstructed spatial representations of visual stimuli at each contrast and under each attention condition using a multivariate inverted encoding model (IEM) applied to visual areas V1-hV4 (see Materials and Methods) (Brouwer and Heeger, 2009b; Sprague and Serences, 2013b; Sprague et al., 2014, 2016, 2018; Vo et al., 2017). Then, we evaluated changes in these spatial reconstructions by quantifying and comparing the amplitude (*A*), size (σ), and reconstruction offset (*B_r_*) measured by fitting a 2-D surface (Figures 2a-b & S2a). An analysis based on 2-way repeated-measures ANOVAs showed that there were significant main effects of contrast (F(5, 30) = 24.63, p < 0.001) and attention (F(1, 6) = 22.00, p = 0.003) but no significant interaction between the two factors on the amplitude parameter of the reconstructions collapsed across V1-hV4 (F(5, 30) = 0.75, p = 0.593), similar to the univariate results. Unlike reconstruction amplitude, there was no main effect of contrast (F(5, 30)’s ≤ 0.96, p’s ≥ 0.458), no main effect of attention (F(1, 6)’s ≤ 0.80, p’s ≥ 0.406), or no interaction between contrast and attention on the size (σ) and reconstruction offset (*B_r_*) of the reconstruction (F(5, 30)’s ≤ 1.62, p’s ≥ 1.86).

We then used the amplitude parameter (A) of the best fitting Gaussian surface to generate a CRF based on the reconstructions. Like the univariate fMRI result, we fit each CRF using a Naka-Rushton equation and found that the CRF baseline parameter (*b_c_*) increased with attention in nearly all visual areas (Figure 2a and S2, resampling tests, p’s < 0.001 for V1, V2/3, and V1-hV4, except that p = 0.310 for hV4, also see Table S2). However, attention effects on the other parameters including *R_max_, C_50_*, and *n* were less robust and less consistent across different visual areas: *p*-values ranged from 0.044-0.946, from 0.311-0.704, and from 0.465-0.753 for *R_max_, C_50_*, and n, respectively (also see Table S2). Taken together, the fMRI analyses provide evidence that attention primarily operates to increase the baseline offset of CRFs based either on univariate responses or on the amplitude of multivariate reconstructions (Figure 1a right), and multivariate assays of information content demonstrate this effect is confined to a change in the signal-to-noise ratio of the stimulus representation (Figure 2a) (Buracas and Boynton, 2007; Murray, 2008; Pestilli et al., 2011; Gouws et al., 2014; Hara and Gardner, 2014; Sprague et al., 2018).

### SSVEP results

In contrast to changes in the CRF baseline activity that were observed in the fMRI data, stimulus-evoked responses measured via EEG revealed a qualitatively different pattern of neural modulations. First, we examined attentional modulations of steady-state visually evoked potentials (SSVEPs), the phase-locked EEG responses in visual cortex that oscillate at the second harmonic (i.e., 30 Hz) of the frequency of the contrast-reversing flickering visual stimulus at 15 Hz (Fig. 3A middle/right) (Kim et al., 2007, 2011; Norcia et al., 2015). For the SSVEP-based CRF, we observed an increase in *R_max_* (response gain) as well as an increase in *C_50_* (a right shift of the CRF) for the attend-stimulus condition compared to the attend-fixation condition (see Figure 3A left, resampling tests, p = 0.048 and p = 0.021 for *R_max_* and *C_50_*, respectively, also see Table S3). However, the *b_c_* and *n* parameters of the SSVEP-based CRFs did not differ across attention conditions (resampling tests, p’s ≥ 0.479; see also Table S3).

**Figure 3.**
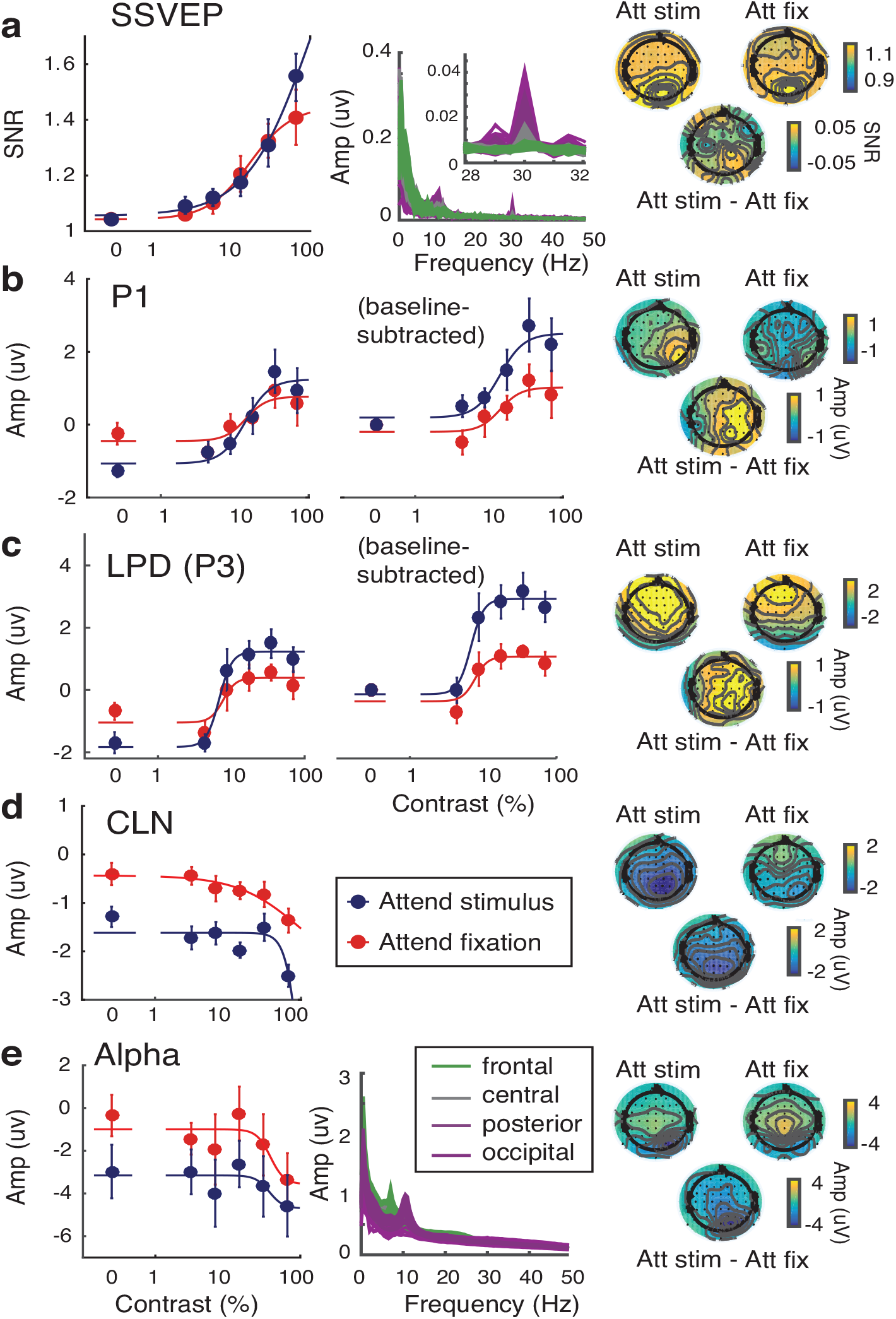
Spatial attention enhances response gain in evoked EEG components and CRF baseline activity in sustained components and induced alpha power. Spatial attention enhanced response gain (i.e., slope) of neural CRFs based on SSVEPs **(a)** and stimulus-evoked ERP components including the contralateral visual P1 **(b)** and the late positive deflection (LPD or P3) **(c)**. In contrast, spatial attention increased the CRF baseline activity of the contralateral late negativity (CLN) **(d)** and post-stimulus alpha amplitude **(e)**. The middle panel in **(a)** demonstrates the frequency plot of the evoked oscillatory EEG signals where SSVEPs were sharply tuned at 30 Hz (i.e., a second harmonic of contrast-reversing stimulus flickering at 15 Hz) and peaked at the occipital electrodes. The middle panels in **(b)** and **(c)** demonstrate the neural CRFs based on the P1 and LPD components where the best fit baseline parameters were subtracted to better illustrate changes in the slopes of the CRFs. The middle panel in **(e)** demonstrates the frequency plot of the induced oscillatory EEG signals where alpha activity peaked at ~12Hz in the occipital and posterior electrodes. The right panels show the topographical maps averaged across all contrast levels in the attend-stimulus and attend-fixation conditions (top left and right, respectively) and the difference between the two conditions (bottom). The left and right sides of the map represent positions ipsilateral and contralateral to the stimulus, respectively. Error bars represent 68% CIs from resampling procedures.

### Early stimulus-evoked ERP results

We also observed qualitatively similar results for early stimulus-evoked event-related potentials (ERPs), including the contralateral occipital P1 component (120-130 ms) and the central-posterior late positive deflection (LPD or P3) (250-350 ms). Specifically, the *R_max_* parameters associated with the P1- and the LPD-based CRFs increased with attention (Figures 3b-c, resampling tests, p’s < 0.001 for both of the P1 and LPD, also see Table S3). Also, the CRF baseline activity (*b_c_*) of these two ERP components were significantly more negative in the attend-stimulus condition than in the attend-fixation condition (resampling tests, p = 0.014 and p = 0.005 for the P1 and LPD components, respectively, also see Table S3). However, note that the direction of these attentional modulations on the CRF baseline parameter was opposite to the polarity of these ERP components and to the direction of contrast modulations, standing in contrast with the fMRI results. We believe this apparent reversal was likely driven by a slow negative-going ERP induced by sustained covert spatial attention (Woodman and Luck, 1999; Vogel and Machizawa, 2004; Vogel et al., 2005; Woodman et al., 2009; Carlisle et al., 2011; Kuo et al., 2012; Tsubomi et al., 2013) (see Figure S3). The other best-fit CRF parameters based on the P1 and LPD components, including the *C_50_* and *n* parameters, did not change (resampling tests, p’s ≥ 0.298 also see Table S3). In addition, while the amplitude of the contralateral visual N1 (150-170 ms) increased as a function of contrast (i.e., became more negative), we observed no attention modulation of any of the CRF parameters (Figure S4, resampling tests, p’s ≥ 0.366, also see Table S3). Overall, the results based on stimulus-evoked responses including SSVEP, P1, and LPD provide converging evidence that attention increases the response gain of neural CRFs (Figure 1a, left).

### Late slow-going ERP results

Discrepancies between fMRI and EEG measures of early stimulus-evoked responses (SSVEP, P1 and LPD/P3) left us wondering whether we could find any similarity in attentional modulations of these two methods. We found that in a much later time window (800-2000 ms), there was an increase in the CRF baseline parameter (*b_c_*) associated with the EEG-based CRF driven by a slow negative-going wave in occipital electrodes that were contralateral to the attended target (Figure 3d, resampling test, p < 0.001, also see Table S3). This modulation had a selective impact on the CRF baseline parameter, as the *R_max_, C_50_* and *n* parameters were not different across attention conditions (resampling tests, p’s ≥ 0.764, also see Table S3). This contralateral slow negative-going wave (termed here the contralateral late negative potential, or CLN) has a characteristic temporal window, polarity, and electrode location that resembles the contralateral delay activity, which is a marker of the active maintenance of attention during visual search and the maintenance of information in working memory (Woodman and Luck, 1999; Vogel and Machizawa, 2004; Vogel et al., 2005; Woodman et al., 2009; Carlisle et al., 2011; Kuo et al., 2012; Tsubomi et al., 2013). The present data show that this ERP component also indexes sustained covert spatial attention even in the absence of a stimulus and that the magnitude of attention modulations was independent of stimulus contrast, consistent with our control over behavioral performance (Figure 1c). This marker thus exhibits a pattern of additive modulation with attention that is similar in a nature to the pattern of BOLD responses.

### Alpha results

We also measured the reduction of contralateral posterior alpha activity in the EEG data (10-12 Hz) relative to pre-cue baseline (Figure 3e). We hypothesized that attentional modulations of alpha oscillations might be similar to that of BOLD activation for two reasons. First, a recent study measured fMRI and electrocorticography (ECoG) and found a strong trial-by-trial correlation between BOLD responses and alpha oscillations in local field potentials recorded from human visual cortex (V1-V3) (Conner et al., 2012; Hermes et al., 2017). Second, alpha oscillations have been shown to track the allocation of selective spatial attention (Foxe et al., 1998; Fries, 2001; Sauseng et al., 2005; Kelly et al., 2006, 2009; Klimesch et al., 2007; Rihs et al., 2007; Fries et al., 2008; Bosman et al., 2012; Foster et al., 2016; Samaha et al., 2016; Foster et al., 2017; Voytek et al., 2017) in a retinotopically selective manner, similar to large-scale patterns of fMRI activity (Kastner et al., 1998, 1999; Gandhi et al., 1999; Sprague and Serences, 2013a; Vo et al., 2017; Sprague et al., 2018). As predicted, we found that CRF baseline activity (*b_c_*) based on post-stimulus alpha amplitude was significantly reduced with attention (resampling test, p = 0.003, also see Table S3). However, the other parameters including *R_max_, C_50_*, and *n* did not differ across attention conditions (resampling tests, p’s ≥ 0.362, also see Table S3). Recently, studies have shown that the topographic patterns of alpha reduction after attention cues and during a working memory delay period contain information about attended and remembered spatial locations even in the absence of continuous sensory input (Sauseng et al., 2005; Kelly et al., 2009; Foxe and Snyder, 2011; Rohenkohl and Nobre, 2011; Bosman et al., 2012; Foster et al., 2016, 2017; Fukuda et al., 2016; Samaha et al., 2016; Green et al., 2017). Our results are consistent with these previous findings and add that attention-induced changes in contralateral alpha power are independent of stimulus contrast, much like the BOLD response.

## Discussion

Overall, the present results demonstrate that BOLD signals in V1-hV4 measured using fMRI and SSVEPs and evoked potentials (P1 and LPD/P3) measured using EEG indexed qualitatively different attentional modulations in the same human subjects performing the same attention task. Notably, visually evoked fMRI responses are most commonly linked with modulations of the SSVEP, P1, and LPD signals, so our demonstration of divergent modulations calls this practice into question. However, we discovered that there were some similarities between the CRFs based on BOLD signals and the CRFs based on the CLN and post-stimulus alpha oscillations. These results suggest that early stimulus-evoked potentials measured using EEG interact with spatial attention, giving rise to response gain modulations in the neural CRFs. However, fMRI signals and other EEG measurements including the contralateral slow-going negativity and post-stimulus alpha activity reflect top-down attentional modulations that are spatially selective but are largely independent of the strength of exogenous stimulus drive.

Consistent with our univariate fMRI results, previous fMRI studies have shown that spatial attention produced an increase in the CRF baseline activity measured via fMRI (Buracas and Boynton, 2007; Murray, 2008; Pestilli et al., 2011; Gouws et al., 2014; Hara and Gardner, 2014; Sprague et al., 2018); only one past fMRI study showed evidence for contrast gain, but the result may have been influenced by different levels of difficulty across stimulus contrast and attention conditions (Li et al., 2008), which we carefully controlled here (Figure 1c). The present study also shows that attention-induced additive shifts in the fMRI-based CRFs were not due to the specific analysis methods applied to the fMRI data because traditional univariate methods that quantify the mean response across all voxels in a visual area and the multivariate IEM yielded similar additive shifts due to attention. In either case, the additive effects of attention on the fMRI-based CRFs are qualitatively different from the response gain that was observed in the SSVEP and ERP measures recorded from the same experimental design and the same participants.

Variants of computational models based on divisive normalization have previously provided at least two alternative explanations for the shift in baseline activity of the fMRI-based CRFs. First, these baseline shifts could be related to the spatial extent of attention and the spatial extent of the stimulus, with additive shifts most prominent when these two factors are similar in size (Herrmann et al., 2012). Alternatively, this attention-related increase in CRF baseline activity could be due to the fact that fMRI signals reflect aggregated neural responses pooled from populations of neurons that exhibit different gain patterns (i.e., contrast or response gain) (Boynton, 2011; Hara et al., 2014), as well as local synaptic activity which can result from long-range projections (Logothetis and Wandell, 2004; Goense and Logothetis, 2008; Magri et al., 2012).

We argue that these explanations, though possible, are not likely to completely account for the observed differences in fMRI and EEG modulations with attention. First, subjects in the present study performed the exact same task using identical visual stimuli and their behavioral performance was equated across measurement modalities. Moreover, fMRI and EEG are both population-level measures that aggregate information across large populations of responsive neurons, yet they still exhibit substantially different patterns of modulation with changes in cognitive demands. This is especially important because recent fMRI and EEG studies using quantitative modeling to link attentional modulations of neural CRFs and psychophysical performance have reached very different conclusions about neural mechanisms underlying attentional selection (Pestilli et al., 2011; Hara and Gardner, 2014; Itthipuripat et al., 2014a, 2017). Specifically, fMRI data suggested that sensory gain (i.e., the increase in CRF slopes) and noise reduction (i.e., the decrease in trial-by-trial variability in neural responses) had limited roles in supporting attention-induced changes in behavioral performance (Pestilli et al., 2011; Hara and Gardner, 2014). In contrast, EEG data suggested that sensory gain modulations and in some cases noise reduction could sufficiently account for attentional benefits in behavior (Mangun and Hillyard, 1990; Störmer et al., 2009; Itthipuripat et al., 2014a, 2017). The results from the present study help to reconcile these different conclusions and point to differences in the physiological sensitivity of neural recording techniques rather than other task and stimulus parameters.

Collectively, our data challenge the general belief that attention-induced changes in fMRI and stimulus-evoked EEG responses merely reflect similar neural processes at different spatial and temporal resolutions. Instead, the findings emphasize that different physiological processes assayed by these complementary techniques must be jointly considered when making inferences about the neural underpinnings of cognitive operations (Boynton, 2011; Hara et al., 2014; Itthipuripat and Serences, 2015) and when using them as diagnostic tools that measure disruptions in cognitive and sensory functions in clinical populations (Calderone et al., 2013).

---

**Figure S1.**
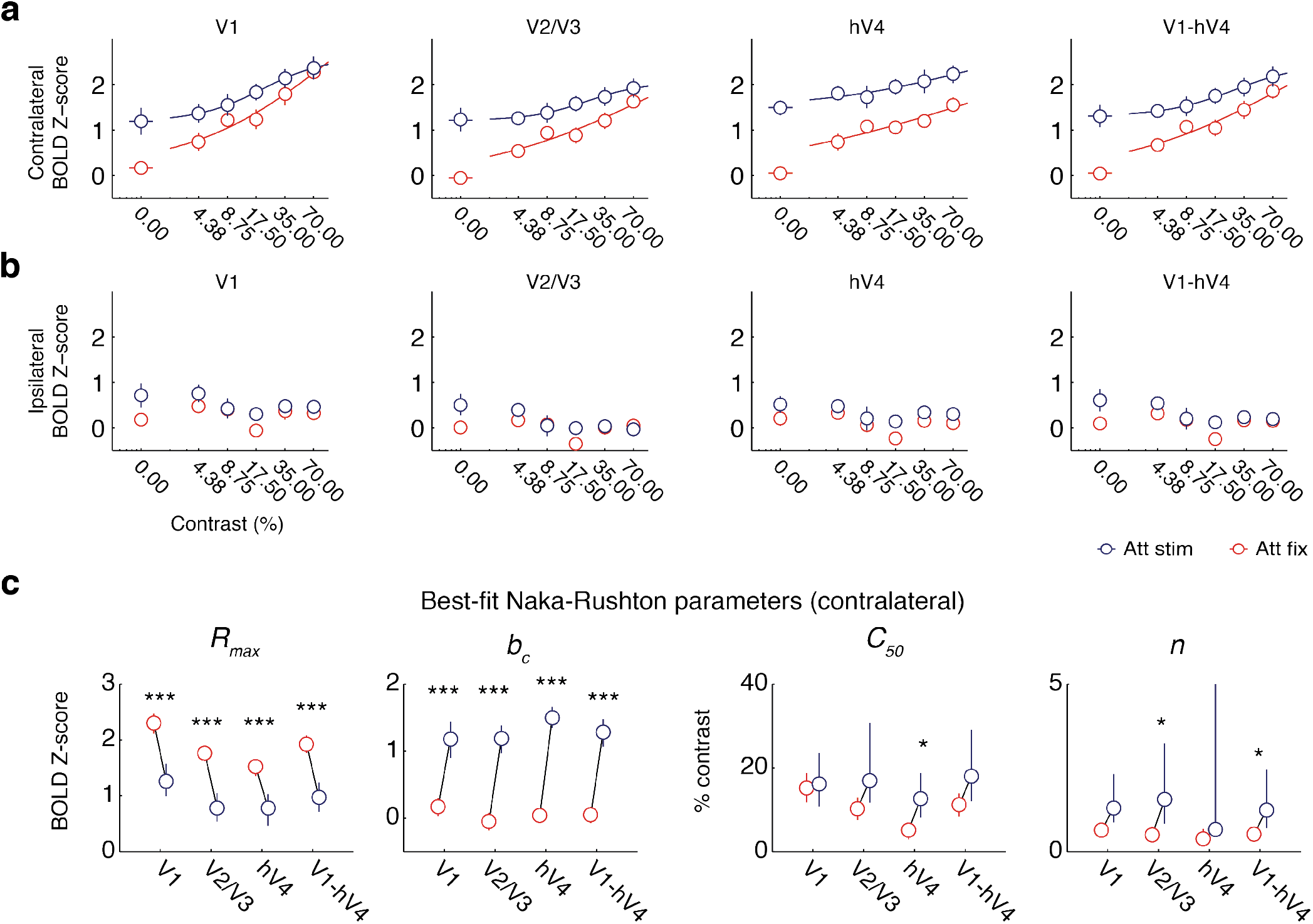
Univariate fMRI results across visual areas V1, V2/3, hV4, and V1-hV4 (see corresponding statistical results in Table S1). **(a)** Attentional modulations of the contrast response functions (CRFs) measured across visual areas contralateral to stimulus presentation. **(b)** Same comparisons for ipsilateral areas. **(c)** Corresponding fit parameters of the CRFs measured from contralateral areas shown in **(a)**. * and *** indicated significant differences between attention conditions (red = attend-fixation and blue = attend-stimulus) with p’s < 0.05 and p’s < 0.001, respectively. Error bars represent 68% CIs from resampling procedures.

**Figure S2.**
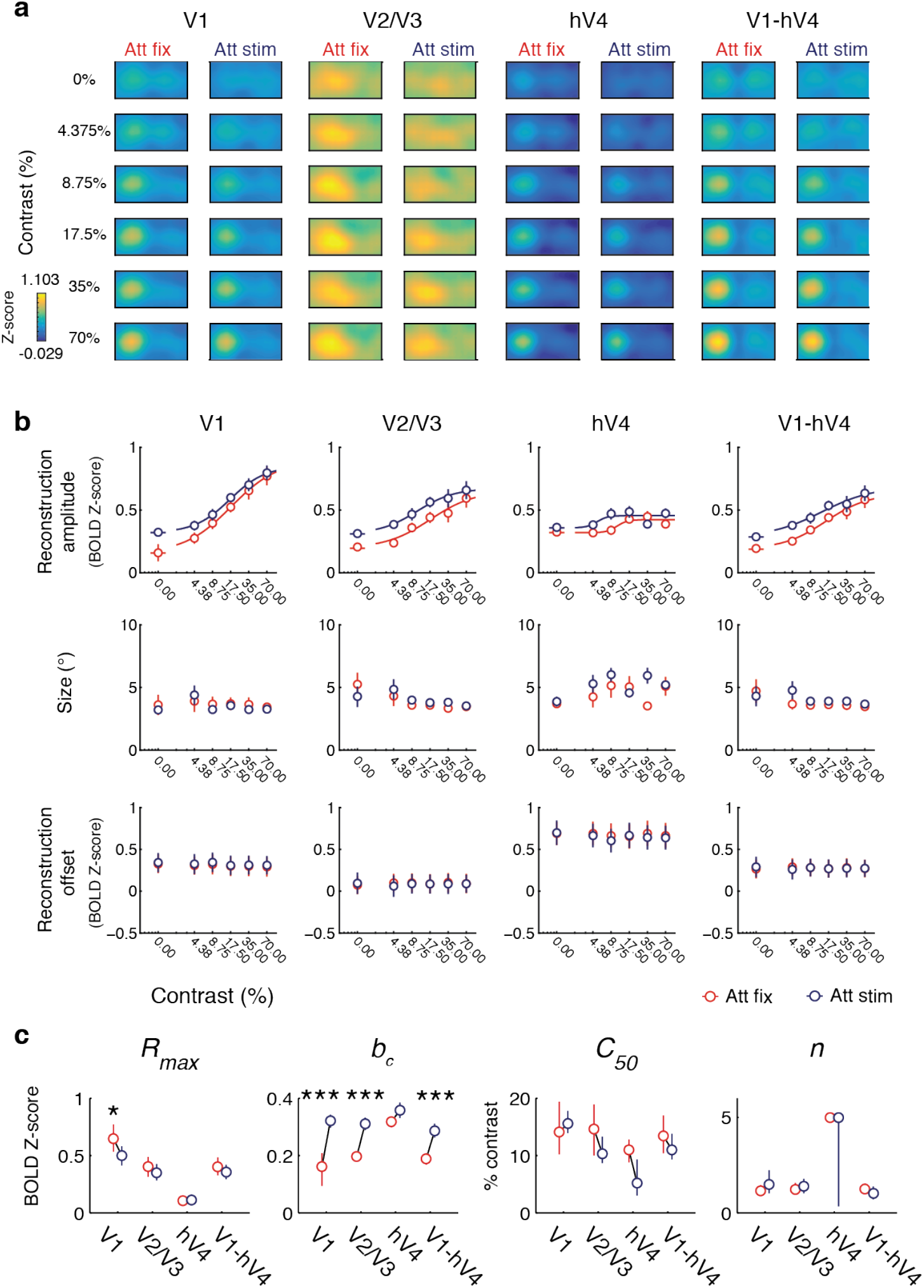
Multivariate fMRI results across visual areas V1, V2/3, hV4, and V1-hV4 (see corresponding statistical results in Table S2). **(a)** The spatial reconstructions of visual stimuli across different contrast and attention conditions. **(b)** The plots of the amplitude, size, and reconstruction offset parameters of the spatial reconstructions shown in **(a)** as a function of stimulus contrast across different attention conditions. **(c)** Corresponding Naka-Rushton fit parameters of the CRFs based on the amplitude (*A*) of surfaces fit to spatial reconstructions, shown in the top panels of **(b)**. * and *** indicated significant differences between attention conditions (red = attend-fixation and blue = attend-stimulus) with *p*’s < 0.05 and p’s < 0.001, respectively. Error bars represent 68% CIs from resampling procedures.

**Figure S3.**
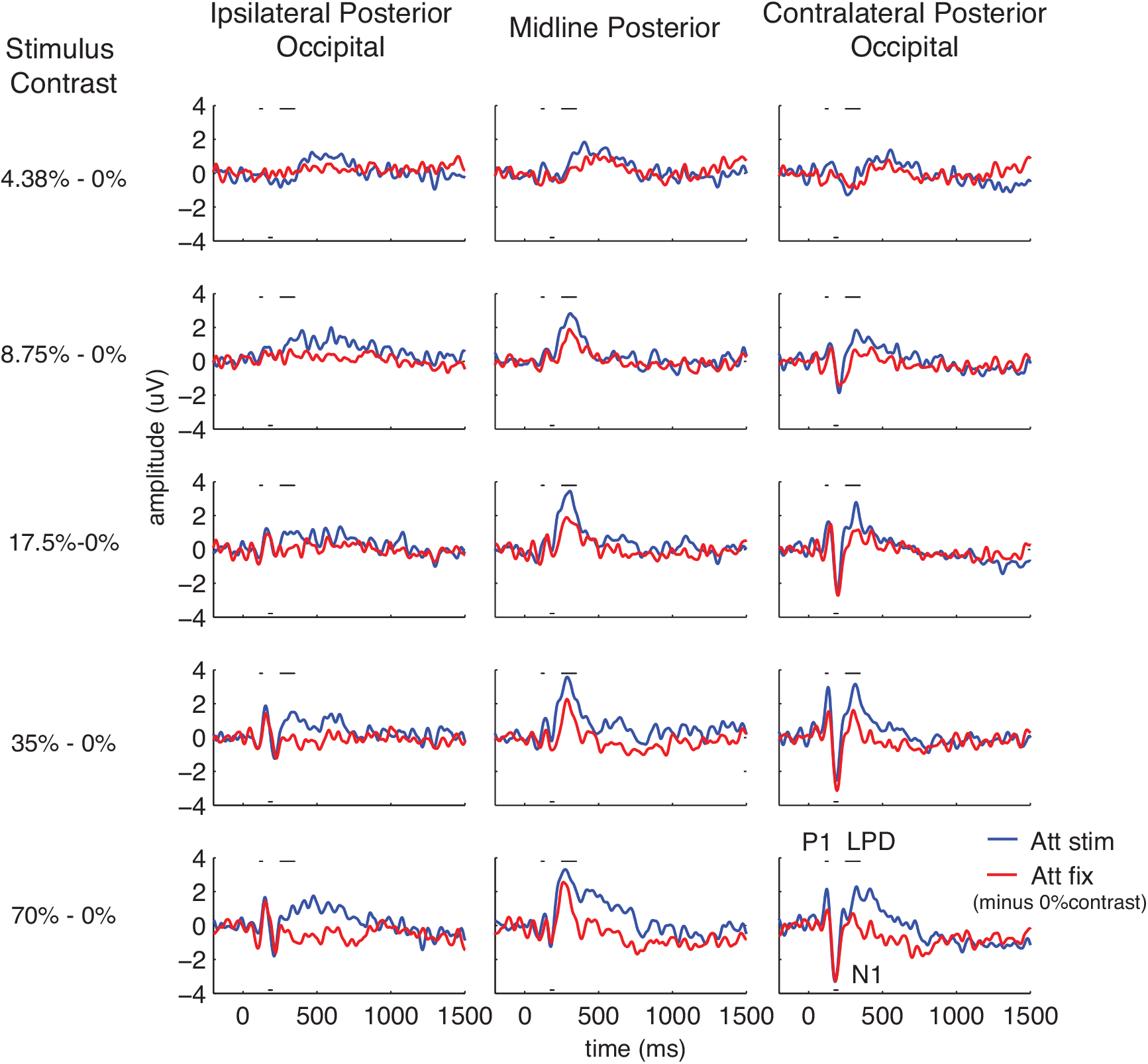
ERP traces evoked by stimuli of different contrast levels across different attention conditions. The ERP trace elicited on stimulus-absent trials (0% contrast) was subtracted from the ERPs evoked by stimuli of all other contrast levels (4.38%-70%) to remove a slow-going negative potential that appeared to have an attention effect that was independent of stimulus contrast (see Figures 3d and S4). The subtracted waveforms contained the visual P1 component, the visual N1 component, and the late positive deflection (LPD or P3) that peaked from 120-130 ms at the contralateral posterior occipital electrodes, from 150-170 ms at the contralateral posterior occipital electrodes, and 250-350 ms at the midline posterior electrodes, respectively.

**Figure S4.**
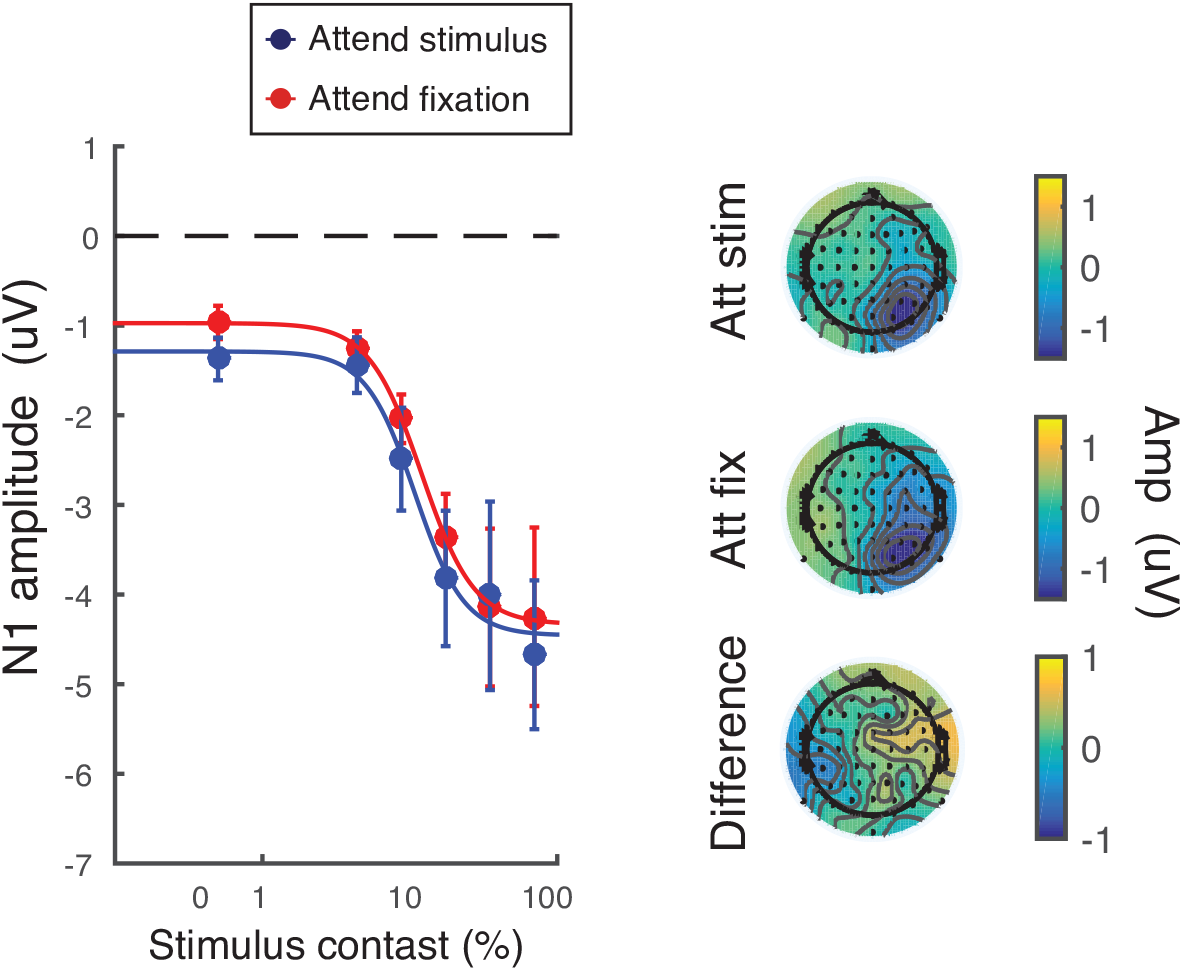
CRFs based on the amplitude of the N1 component. Topographical maps show the data collapsed across all contrast levels. Left and right sides of the maps represent positions ipsilateral and contralateral to the stimulus, respectively. Error bars represent 68% CIs from resampling procedures.

**Figure S5.**
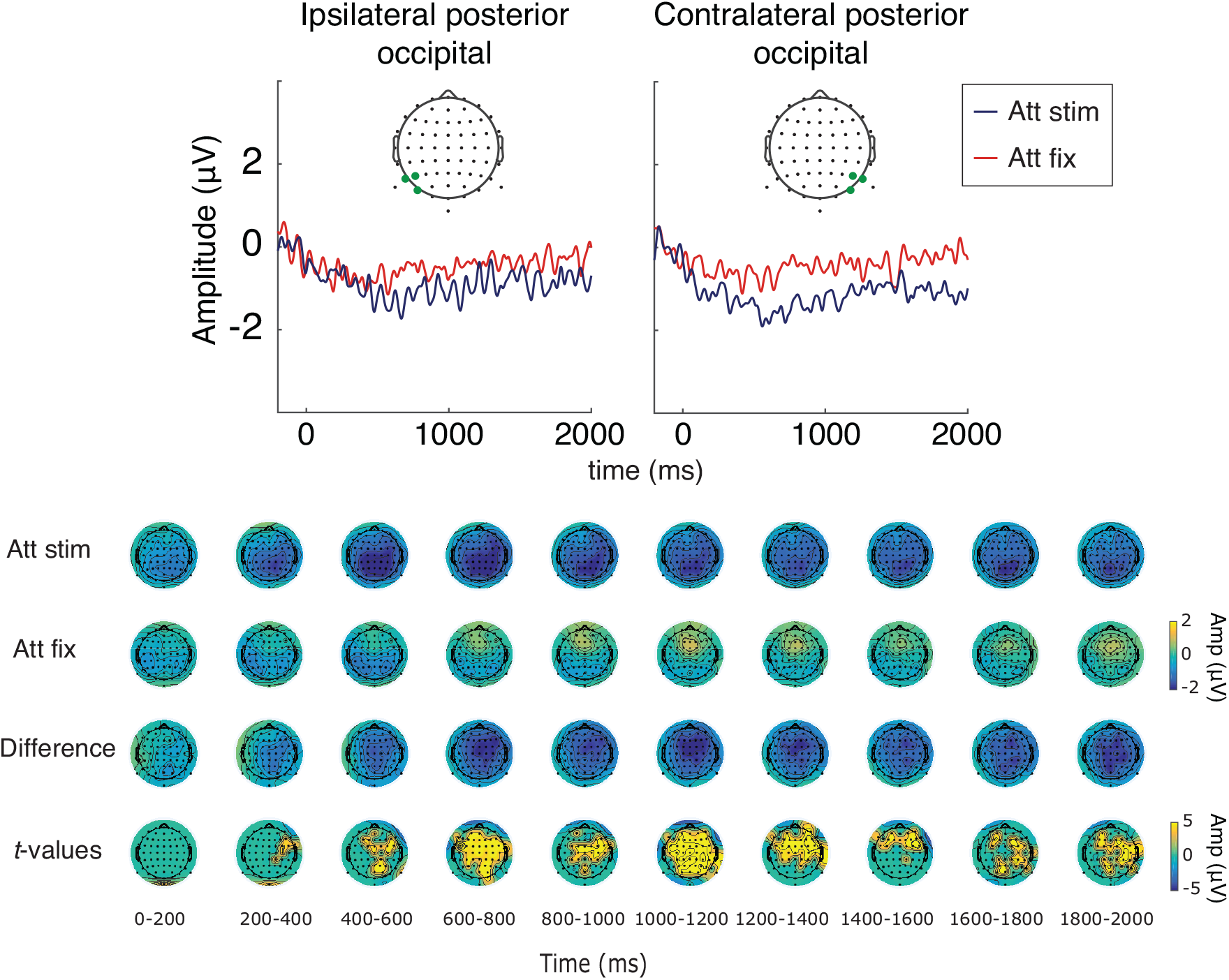
The slow-going negative waveform measured in the contralateral posterior occipital electrodes, termed here as contralateral late negativity (CNL). This ERP component increased in amplitude when attention was directed to the stimulus location even in the absence of a visual stimulus (stimulus contrast of 0%). The topographical maps show the data from 0-2000 ms in 10 steps of 200 ms. The T-values were thresholded to present only the data that passed a significance level of α ≤ 0.05, FDR-corrected (the bottom-most panels). The left and right sides of the maps represent positions ipsilateral and contralateral to the stimulus, respectively.

**Table S1.** Statistical resampling results for univariate fMRI data. (+) and (−) indicate changes in the same and opposite directions as the main effect of contrast on the CRF.

| Univariate fMRI data | Figure | R<sub>max</sub> | C<sub>50</sub> | b<sub>c</sub> | n |
| --- | --- | --- | --- | --- | --- |
|  |  | Attend-fixation[68%CI]<br>Attend-stimulus [68%CI]<br>p-value |  |  |  |
| Contralateral V1 | S1a and S1c | [2.12 1.47]<br>[1.00 1.57]<br>p < 0.001***(-) | [11.80 18.70]<br>[10.80 23.50]<br>p = 0.914 | [0.03 0.30]<br>[0.90 1.44]<br>p < 0.001***(+) | [0.55 0.84]<br>[0.88 2.31]<br>p = 0.084 |
| Contralateral V2/3 | S1a and S1c | [1.62 1.90]<br>[0.54 1.05]<br>p < 0.001***(-) | [7.60 12.90]<br>[11.70 30.70]<br>p = 0.095 | [-0.18 0.08]<br>1.00 1.38]<br>p < 0.001***(+) | [0.43 0.58]<br>[0.84 3.23]<br>p = 0.025* |
| Contralateral hV4 | S1a and S1c | [1.36 1.65]<br>[0.46 1.03]<br>p < 0.001***(-) | [2.90 6.80]<br>[8.10 18.70]<br>p = 0.026*(+) | [-0.05 0.15]<br>[1.35 1.66]<br>p < 0.001***(+) | [0.31 0.68]<br>[0.48 5.00]<br>p = 0.112 |
| Contralateral V1-hV4 | 2a, S1a, and S1c | [1.77 2.08]<br>[0.72 1.23]<br>p < 0.001***(-) | [8.40 13.90]<br>[12.00 29.10]<br>p = 0.104 | [-0.07 0.17]<br>[1.07 1.48]<br>p < 0.001***(+) | [0.45 0.64]<br>[0.70 2.45]<br>p = 0.040* |

**Table S2.** Statistical resampling results for spatial reconstruction amplitude derived from fMRI data. (+) and (−) indicate changes in the same and opposite directions as the main effect of contrast on the CRF.

| Amplitude<br>(A) of fMRI<br>reconstructi<br>on | Fig. | R<sub>max</sub> | C<sub>50</sub> | b<sub>c</sub> | n |
| --- | --- | --- | --- | --- | --- |
|  |  | Attend-fixation[68%CI] |  |  |  |
|  |  | Attend-stimulus [68%CI] |  |  |  |
|  |  | p-value |  |  |  |
| V1 | S2b-c | [0.53 0.77]<br>[0.41 0.58]<br>p = 0.044*(-) | [10.20 19.40]<br>[13.90 17.80]<br>p = 0.704 | [0.09 0.21]<br>[0.30 0.34]<br>p < 0.001***(+) | [0.88 1.46]<br>[1.02 2.23]<br>p = 0.584 |
| V2/3 | S2b-c | [0.31 0.49]<br>[0.28 0.43]<br>p = 0.282 | [10.00 18.90]<br>[8.70 13.30]<br>p = 0.519 | [0.18 0.21]<br>[0.29 0.33]<br>p < 0.001***(+) | [1.04 1.57]<br>[1.02 1.79]<br>p = 0.753 |
| hV4 | S2b-c | [0.09 0.13]<br>[0.06 0.16]<br>p = 0.946 | [8.80 12.80]<br>[3.00 9.30]<br>p = 0.311 | [0.30 0.34]<br>[0.33 0.38]<br>p = 0.310 | [5.00 5.00]<br>[0.35 5.00]<br>p = 0.465 |
| V1-hV4 | 2d<br>and<br>S2b-c | [0.33 0.48]<br>[0.29 0.42]<br>p = 0.395 | [10.4 17.00]<br>[9.30 13.80]<br>p = 0.603 | [0.17 0.21]<br>[0.26 0.31]<br>p < 0.001***(+) | [1.01 1.55]<br>[0.70 1.38]<br>p = 0.474 |

**Table S3.** Statistical resampling results for all EEG measurements. (+) and (−) indicate changes in the same and opposite directions as the main effect of contrast on the CRF.

| EEG Measurements | Figure | R<sub>max</sub> | C<sub>50</sub> | b<sub>c</sub> | n |
| --- | --- | --- | --- | --- | --- |
|  |  | Attend-fixation[68%CI]<br>Attend-stimulus [68%CI]<br>p-value |  |  |  |
| SSVEP | 3a | [0.28 0.52]<br>[0.47 0.78]<br>p = 0.048*(+) | [14.60 33.00]<br>[27.00 54.00]<br>p = 0.021*(-) | [1.03 1.06]<br>[1.04 1.08]<br>p = 0.479 | [1.39 3.63]<br>[1.20 2.20]<br>p = 0.539 |
| P1 | 3b | [0.72 1.83]<br>[1.67 3.01]<br>p < 0.001***(+) | [7.20 18.60]<br>[11.90 17.50]<br>p = 0.797 | [-0.70 -0.30]<br>[-1.20 -0.93]<br>p = 0.014*(-) | [2.74 5.00]<br>[1.89 5.00]<br>p = 0.372 |
| N1 | S3 | [-4.39 -2.27]<br>[-4.28 -2.31]<br>p = 0.826 | [9.50 14.10]<br>[9.60 14.60]<br>p = 0.811 | [-1.14 -0.84]<br>[-1.48 -1.08]<br>p = 0.366 | [2.04 4.14]<br>[1.49 5.00]<br>p = 0.4906 |
| LPD | 3c | [1.23 1.73]<br>[2.66 3.55]<br>p < 0.001***(+) | [6.60 10.40]<br>[6.00 7.80]<br>p = 0.298 | [-1.33 -0.77]<br>[-2.11 -1.60]<br>p = 0.005**(-) | [5 5]<br>[5 5]<br>p = 0.723 |
| CNL | 3d | [-1.57 -0.90]<br>[-2.98 -0.62]<br>p = 0.764 | [15.70 62.70]<br>[3.10 80.3]<br>p = 0.863 | [-0.63 -0.19]<br>[-1.63 -1.13]<br>p < 0.001***(+) | [0.68 4.03]<br>[0.32 5.00]<br>p = 0.769 |
| Alpha | 3e | [-4.81 -2.07]<br>[-3.26 -0.86]<br>p = 0.362 | [6.50 63.20]<br>[12.70 60.40]<br>p = 0.885 | [-1.61 0.36]<br>[-4.22 -1.75]<br>p = 0.003**(+) | [0.46 5.00]<br>[0.81 5.00]<br>p = 0.719 |

## Acknowledgments & author contributions

SI and TCS conceived, implemented the experiments, collected and analyzed the data, and cowrote the manuscript. JTS conceived and supervised the project and co-wrote the manuscript. Funding was provided by NIH R01-EY025872 (J.T.S.), the James S. McDonnell Foundation (J.T.S), the Howard Hughes Medical Institute International program (S.I.), a Royal Thai Scholarship from the Ministry of Science and Technology in Thailand (S.I.), NIH T32-MH020002 (T.C.S.), NIH T32-EY007136 (T.C.S.), and NIH F32-EY023438 (T.C.S.).

